# Employing a honeybee olfactory neural circuit as a novel gas sensor for the detection of human lung cancer biomarkers

**DOI:** 10.1101/2023.10.04.560899

**Authors:** Michael Parnas, Elyssa Cox, Simon Sanchez, Alexander Farnum, Noël Lefevre, Sydney Miller, Debajit Saha

## Abstract

Human breath contains biomarkers (odorants) that can be targeted for early disease detection. It is well known that honeybees have a keen sense of smell and can detect a wide variety of odors at low concentrations. Here, for the first time, we employ honeybee olfactory neuronal circuitry to classify human lung cancer volatile biomarkers and their mixtures at concentration ranges relevant to human breath, parts-per-billion to parts-per-trillion. Different lung cancer biomarkers evoked distinct spiking response dynamics in the honeybee antennal lobe neurons indicating that those neurons encoded biomarker-specific information. By investigating lung cancer biomarker-evoked population neuronal responses from the honeybee antennal lobe, we could classify individual human lung cancer biomarkers successfully (88% success rate). When we mixed six lung cancer biomarkers at different concentrations to create ‘synthetic lung cancer’ vs. ‘synthetic healthy breath’, honeybee population neuronal responses were also able to classify those complex breath mixtures successfully (100% success rate with a leave-one-trial-out method). Finally, we used separate training and testing datasets containing responses to the synthetic lung cancer and healthy breath mixtures. We identified a simple metric, the peak response of the neuronal ensemble, with the ability to distinguish synthetic lung cancer breath from the healthy breath with 86.7% success rate. This study provides proof-of-concept results that a powerful biological gas sensor, the honeybee olfactory system, can be used to detect human lung cancer biomarkers and their complex mixtures at biological concentrations.

## 1. Introduction

Honeybees have a sensitive olfactory system designed to help them navigate complex environments encompassing foraging, reproduction, brood care, and defense. Consequently, they can be used to reliably detect a wide range of volatile chemicals, or the ‘smell’ of objects, while also distinguishing between odor mixtures efficiently even at low concentrations^1–4^. Here, we hypothesize that the honeybee’s powerful olfactory neural circuitry can be leveraged to develop a gas sensing system with the ability to detect lung cancer biomarkers present in exhaled human breath.

Lung cancer is the second most commonly diagnosed cancer worldwide, and is the leading cause of cancer-related death among both men and women^5^. Reliable and early detection of lung cancer can improve patient outcomes. The analysis of volatile organic compounds (VOCs) in exhaled human breath is a promising avenue for detecting metabolically linked diseases such as cancer at an early stage^6–15^. Therefore, breath-based VOC analysis can potentially answer the critical need for sensitive, early, and noninvasive cancer diagnostics^16,17^. Exhaled human breath contains over 3,500 known VOCs^16,18,19^. Different diseases alter the components and concentrations of these VOCs, and thus can reflect the metabolic condition, or health, of an individual^20–24^. Several studies employing exhaled human breath and mass spectrometry (e.g., GC-MS) have shown that cancer can alter specific VOCs in exhaled breath at parts-per-billion (ppb) to parts-per-trillion (ppt) ranges^7,17^. In this study, we focus on VOCs that are altered in patients with lung cancer.

Generally, VOC analysis from exhaled breath is performed using mass spectrometry-based analysis techniques (such as GC-MS, SIFT-MS, PTR-MS, and GC-IMS)^6,9,21,25–46^. Although, GC-MS, the current gold standard, has high sensitivity and specificity when identifying known compounds, it has difficulty in identifying correct concentrations of unknown compounds at ppb to ppt levels and requires pre- and post-processing of data, which is not standardized. Another technology for VOC sensing is electronic noses (e-noses) which employ some biological principles for one-shot gas sensing. These e-nose devices are portable and can detect a few target compounds at low (ppb) concentrations. E-nose devices have been successfully employed to detect multiple types of cancer^14,15,30,47–60^. However, e-noses are usually targeted for a specific chemical, and these engineered sensors cannot match the broad sensitivity and specificity of biological chemical sensors^61–67^.

Biological “noses”, such as the honeybee antennae and olfactory brain are extremely sensitive, and honeybees have been shown to learn odor identity and perform complex olfactory behavioral tasks^1,3,68^. Similar to dogs’ noses which have been successful in the detection of different VOCs of interests^62,69–75^, it has been demonstrated that insects’ noses, or antennae, can be exposed to a target smell and reinforced with a food reward for detection of that target stimuli, behaviorally^76^. Here, we take advantage of the honeybees’ sensitive olfactory system to test whether the central olfactory neural circuitry (i.e., the antennal lobe) can generate discriminatory neural responses to different human lung cancer biomarkers and their mixtures. In the honeybee olfactory sensory pathway, olfactory receptor neurons (ORNs) located in the antenna convert chemical cues into electrical signals. Signals from ORNs are carried to the antennal lobe where two types of neurons, projection neurons (PNs) and local neurons (LNs), form dense clusters of connections called glomeruli. The roughly 60,000 ORNs present in the honeybee antenna send their output to only 800 PNs in the antennal lobe^77^, which transmit odor-evoked spiking responses to 368,000 Kenyon cells in the mushroom body^78^, making the antennal lobe an ideal location for neural recordings due to the convergence of VOC-evoked spiking information.

In this study, we have systematically tested two hypotheses: (1) honeybee antennal lobe neurons (ALNs) possess discriminatory information corresponding to multiple human lung cancer biomarkers, and (2) these neurons can differentiate between small differences in concentrations of the lung cancer biomarker mixtures (i.e., synthetic lung cancer vs. synthetic healthy breath mixtures). To achieve these goals, we have performed *in vivo* electrophysiological recordings from the honeybee antennal lobe while exposing the honeybee antennae to lung cancer biomarkers and mixtures. We analyzed the lung cancer VOC-evoked neural data using biological neural computational schemes to classify lung cancer biomarkers and to test our hypotheses.

## 2. Results

### 2.1. Human lung cancer biomarkers are detected by the neurons in the honeybee antennal lobe

We began our investigation by obtaining individual lung cancer VOC-evoked neural responses from the honeybee antennal lobe circuit. The rational for targeting antennal lobe circuitry was twofold: (1) since 60,000 ORNs from the antennae converge to only 800 PNs in the honeybee antennal lobe, the probability of getting odor evoked responses to diverse VOCs is higher from the ALN recordings, and (2) in the honeybee antennal lobe, spatiotemporal response properties of ALNs are odor specific^79,80^. We chose nine different VOCs which are implicated as human exhaled breath lung cancer biomarkers^81,82^. For these sets of experiments, the primary goal was to detect the identity of the volatile chemicals using honeybee neural recordings, therefore, we tested all nine VOCs at a fixed concentration (1% vol/vol, diluted in mineral oil).

The timing and volume of the VOC delivery to the honeybee antennae was carefully controlled while simultaneously conducting *in vivo* extracellular recordings from the ALNs, which contained both PN and LN responses (**Fig. 1a**). We observed that VOC-evoked neural voltage traces from a single recording site can display different spiking response properties to each VOC tested (**Fig. 1b**). Next, we identified individual ALNs by spike sorting (see Experimental Section) and plotted peri-stimulus time histograms and raster plots of individual neurons (total *n =* 44 ALNs). A representative neuron’s odor-evoked responses exhibited distinct differences between all nine VOCs and the mineral oil control (**Fig. 1c**). Not only can the honeybees detect each of the odors, as shown by the odor-evoked responses, we observed that the ALNs also respond to each odor differently.

**Figure 1:**
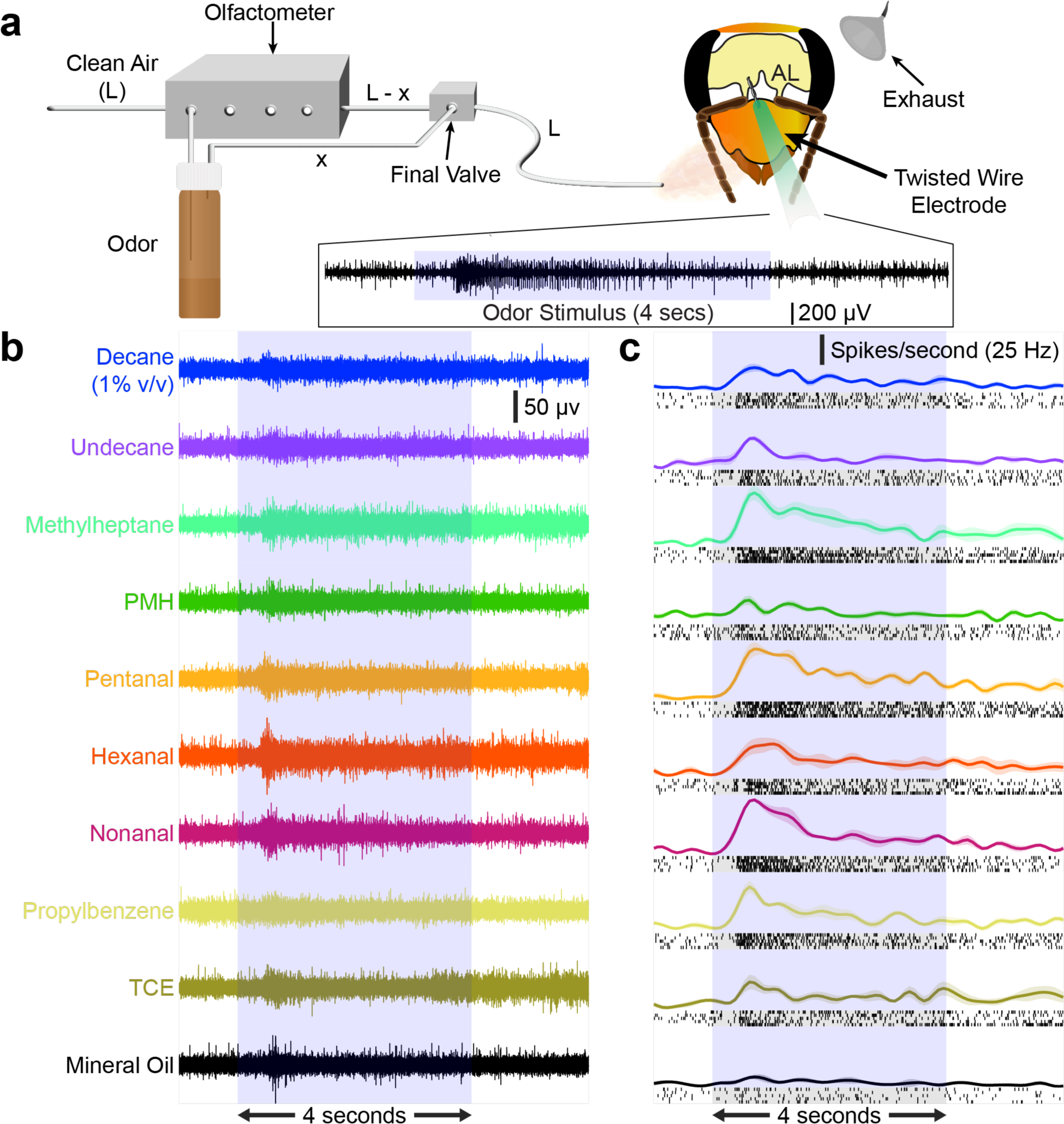
Individual neurons display distinct odor evoked responses to putative lung cancer biomarkers. **(a)** Schematic of the experimental setup. Starting from the left, clean air from a compressed air cylinder enters an olfactometer with a flow rate, L. Prior to odor stimulus, the entire flow rate, L, is delivered directly to the honeybee’s antennae. When the odor stimulus starts, the olfactometer diverts a portion of the air, x, into the headspace of an odor vial that contains a single VOC mixed with mineral oil (1% v/v). The clean air (L-x) and the odor laden air (x) are combined at the final valve and delivered to the antennae. A custom-built twisted wire electrode placed within the honeybee antennal lobe records the VOC-evoked neural responses (shown at the bottom). **(b)** Representative extracellular lung cancer VOC-evoked neural voltage responses from a recording location within the honeybee antennal lobe is shown. Responses to nine putative cancer biomarkers mixed in mineral oil (1% v/v) as well as a pure mineral oil control-evoked response are shown. The light blue box indicates the odor presentation window (4 seconds). **(c)** The representative VOC-evoked voltage responses shown in **panel b** were spike sorted and the raster and peri-stimulus time histograms (PSTHs) of one antennal lobe neuron’s responses to all 10 odors are shown. Each black line in the raster plot indicates an action potential, or spiking event, from the neuron. Raster plots are shown for all five trials of each VOC presentation. The PSTHs (in color) show the changes in spiking rate over time for the neuron with an increase in firing rate correlated to an increase in density of the raster plots. Trial-averaged PSTHs are plotted with the shaded region indicating the S.E.M. The light blue box in the background indicates the 4 second odor presentation window.

### 2.2. Antennal lobe neuron encoding of lung cancer biomarkers are distinct at the population level

Next, we sought to confirm that each ALN contains discriminatory information about odor identity and that the population of ALNs encodes the odors distinctly. To investigate each individual ALN, we first calculated pairwise distances between all 10 odor-evoked responses and averaged them together for each of the 44 neurons (see Experimental Section, **Supplementary Fig. 1**). Notice that a nonzero pairwise distance plot for a neuron indicates that neuron contains discriminatory information for all stimuli tested. The pairwise distance plots showed that the largest separation of VOC-evoked responses for most neurons occurred during the transient phase of the neural response, which lasts about one second from the odor onset (**Fig. 2a**)^83^. We chose a time window from 0.25 – 0.75 seconds (total 500 ms duration) for data analysis as we found that most neurons responded with a 0.25 second delay from the final olfactometer valve opening. This analysis confirmed that all the neurons in the recorded population contained VOC-specific information which can be used to classify lung cancer VOC identity.

**Figure 2:**
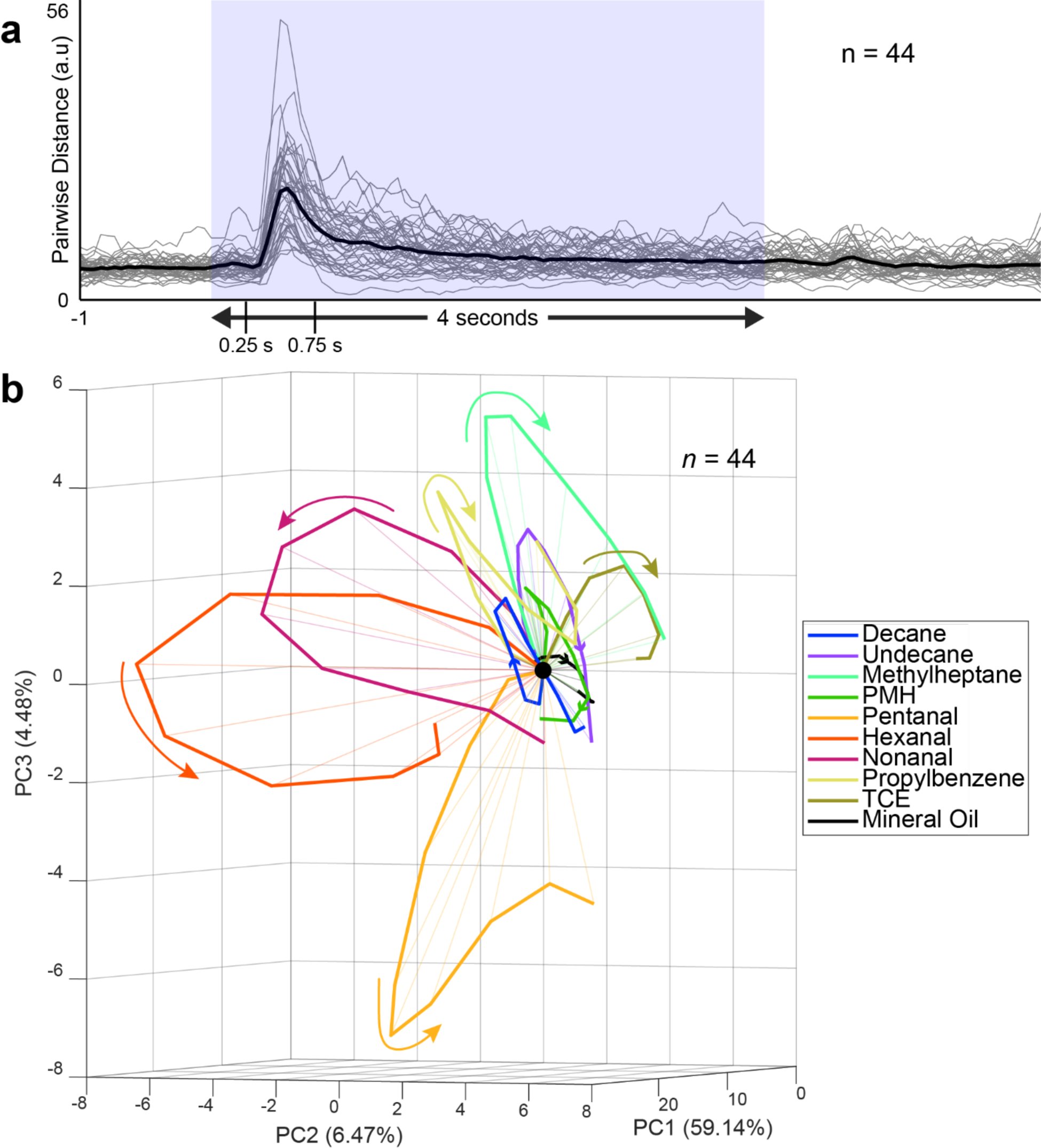
Neural encoding of putative cancer biomarkers is the most distinct during the transient response period. **(a)** Pairwise distance plots (see Experimental Section) are shown for all recorded neurons (n = 44 ALNs total). Each grey line is the average of all possible pairwise distances from a single neuron’s responses to all 10 odors. The black line is the average value of pairwise distances across all 44 neurons. Notice that the average pairwise distances of most neurons have a maximum value during the transient phase of odor stimulus from 0.25 – 0.75 seconds after stimulus onset. **(b)** VOC-evoked population neural trajectories of all 10 odors are shown using PCA dimensionality reduction. The trajectories are plotted between 0.25 – 0.75 seconds after stimulus onset to highlight the transient response window, which is the most discriminatory segment of the neural response. The colored arrows indicate the direction of evolution for each VOC-evoked trajectory. All trajectories are aligned at 0.25 seconds after odor onset (indicated by a black dot), which corresponds to the time of the odor plume hitting the antennae after the opening of the final valve (at 0.0 seconds).

Next, we analyzed the lung cancer biomarker discrimination ability of the entire recorded population of ALNs. To achieve this, multiple neural recordings across honeybees were combined (*n =* 44 ALNs) to generate a lung cancer VOC-evoked population response matrix (*neuron* × *time*). Odor-evoked responses from every neuron were aligned to the odor onset, trial averaged (*n =* 5 trials) firing rates of each neuron corresponding to each odor were binned into 50 ms nonoverlapping bins and then combined to form time-variant, odor-specific, high dimensional population neuronal response vectors. The 50 ms time window selection corresponds to 20 Hz oscillations of odor-evoked local field potentials observed in insect antennal lobes^84–86^.

To visualize the lung cancer VOC-evoked population neuronal trajectories, a principal component analysis (PCA, see Experimental Section) was performed on the high dimensional neural response dataset. For this analysis, a 500 ms time window was chosen from 0.25 – 0.75 second post-stimulus onset, as the ALNs contained the highest amount of discriminatory information about the odor stimuli during this period (**Fig. 2a**). PCA was applied on the entire dataset containing all 10 VOCs and the first three principal components with the highest variance were chosen to project the data onto three dimensions for visualization purposes (**Fig. 2b**). In this reduced space, the odor-evoked population vectors corresponding to each 50 ms time bin were connected in a temporal order to create the neural trajectories for each stimulus and temporally aligned with each other at the origin. Using this analytical technique, previous studies have demonstrated that distinct neural trajectories represent VOC identities and intensities^87^. Here, we observed that neural trajectories corresponding to different VOCs traced different paths in the principal component space, validating that the identity of each lung cancer VOC was uniquely encoded by the honeybee ALN population responses.

### 2.3. Odor-evoked spatiotemporal antennal lobe neuron responses can be leveraged to identify lung cancer biomarkers

We then attempted to quantify the ability of the honeybee ALNs to classify cancer biomarkers by implementing a high-dimensional, leave-one-trial-out (LOTO) analysis scheme (**Figs. 3a-f**, see Experimental Section). First, a high-dimensional population response matrix was constructed as described before (**Figs. 3a, b**). Then, one trial was separated from the neural response matrices for each odor and used as a ‘testing template’ while the remaining four trials were averaged together and used to create the ‘training templates’ (**Fig. 3c**). This resulted in a total of 10 training and 10 testing templates corresponding to nine lung cancer VOCs and the mineral oil control. Time-matched responses within a 50 ms bin of the training and testing templates were plotted in the high dimensional space and the Euclidean distances between each testing template and all 10 training templates were calculated. Testing templates were classified as belonging to the odor whose training template minimized the Euclidean distance (**Fig. 3d**). This LOTO analysis was performed in such a way that each trial (out of the total five trials corresponding to one VOC) becomes the testing template once. Similar to trajectory analysis, this high-dimensional classification was also performed within the 0.25 – 0.75 second post-stimulus onset window. The results are summarized in a confusion matrix (**Figs. 3e, g**), with the testing templates along the X-axis and the training templates along the Y-axis. The higher classification success rates along the diagonal, shown by the darker colors, were indicative of testing templates being classified to the correct training template, resulting in an overall 50.2% success rate. We would expect that completely random assignment of odors would only yield 10% accurate classification in this case. This classification rate reflects how many of the 50 ms population response vectors can be classified correctly. To convert that number to classification success over the entire 500 ms discriminatory neural response window, we took the mode of the 50 ms bin-wise classification values over the duration of each 500 ms trial in a winner-take-all manner (**Figs. 3f, h**). Using this method, we achieved 88% classification success of all cancer biomarkers tested.

**Figure 3:**
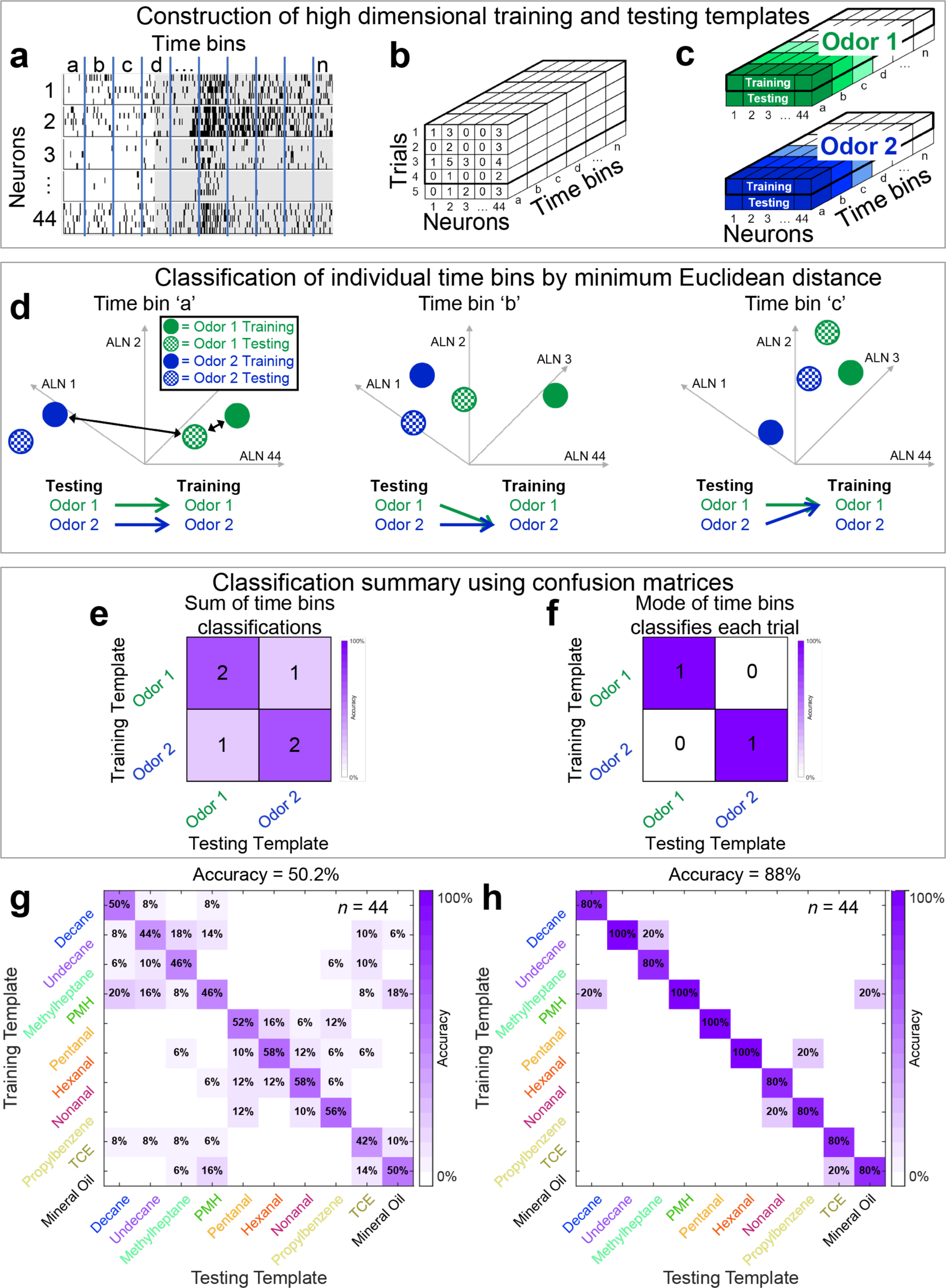
Population neural responses can classify individual lung cancer biomarkers. **(a)** Raster plots of spike sorted neuron responses (1-44) to a single odor (hexanal) from antennal lobe recordings in honeybees. Each neuron’s response was recorded for five trials. The responses are all aligned using the stimulus onset as the reference point and binned into discrete, non-overlapping, 50 ms time bins. **(b)** The number of spiking events in each time bin is counted and used to populate a three-dimensional matrix (neurons x trials x time bins). **(c)** Four trials are selected and averaged together to create the training template, with the fifth trial being *left out* to create the testing template, known as a leave-one-trial-out (LOTO) analysis. Each time bin, denoted by the different shades of color, is analyzed separately. **(d)** The training and testing templates for each time bin and both odors, solid and checkered circles, respectively, are visualized as high dimensional points, with the number of dimensions equal to the number of neurons (n = 44). Testing templates (checkered) are compared to each training template (solid) and assigned using the smallest Euclidean distance. For *time bin a* (left) the green testing template would be assigned to green, and the blue testing template would be assigned to blue. *Time bin b*, (middle) green is assigned to blue and blue is assigned to blue. *Time bin c*, (right) green is assigned to green and blue is assigned to green. **(e)** Using the previous assignments, a confusion matrix can be populated. The testing templates for each odor are on the x-axis and their assignments are on the y-axis. For two time bins (‘a’ and ‘c’), the green testing template was assigned to green, however for one time bin (‘b’) green was mis-assigned to blue. We count the assignments for each of the odors. **(f)** Instead of counting each individual bin, the mode of the bin assignments can be used to assign an entire trial. For the left-out trial visualized here, the mode for the green testing template is green (two out of three) and the mode for the blue testing template is blue (two out of three). After completing the analysis for a single left out trial, we can iterate until each trial has been left out once and used as a testing template. Each time using the other four trials to create the training template. And, this can be expanded to include any number of odors. **(g)** The count of time bin assignments for all nine VOCs and the mineral oil control shows good classification; notice that the diagonal elements of the confusion matrix show higher values compared to the off-diagonal elements which indicates that testing templates are correctly classified in most cases using this quantitative analysis. Overall success rate of classification for this approach is 50.2%. **(h)** Instead of classifying each 50 ms bin individually, here, the entire time window between 0.25 – 0.75 seconds after stimulus onset is being classified in a winner-take-all approach. The mode of the classification value of the 10-time bins (each bin is 50 ms in duration over a total time of 500 ms) was used to determine the overall classification success for the entire testing trial. Using this approach, 44 of the 50 trials were correctly classified with an accuracy of 88%.

### 2.4. Synthetic human lung cancer breath vs. synthetic healthy breath can be differentiated by the honeybee antennal lobe neuronal responses

After verifying that several human lung cancer biomarkers can be detected by the honeybee ALNs (**Figs. 1-3**), we tested if complex mixtures that replicate actual concentrations of these VOCs in healthy individuals’ vs. lung cancer patients’ exhaled breath can be detected by the honeybee ALN responses. We defined these complex VOC mixtures as ‘Synthetic breath’ mixtures (**Fig. 4a**). We leveraged previously published concentration data of the VOCs^7,88^, combined six VOCs at specific concentrations as found in lung cancer vs. healthy human exhaled breath samples, and mixed them in clean air (80% nitrogen and 20% oxygen) to reach the desired biological concentrations at ppb to ppt ranges (**Fig. 4b**).

**Figure 4:**
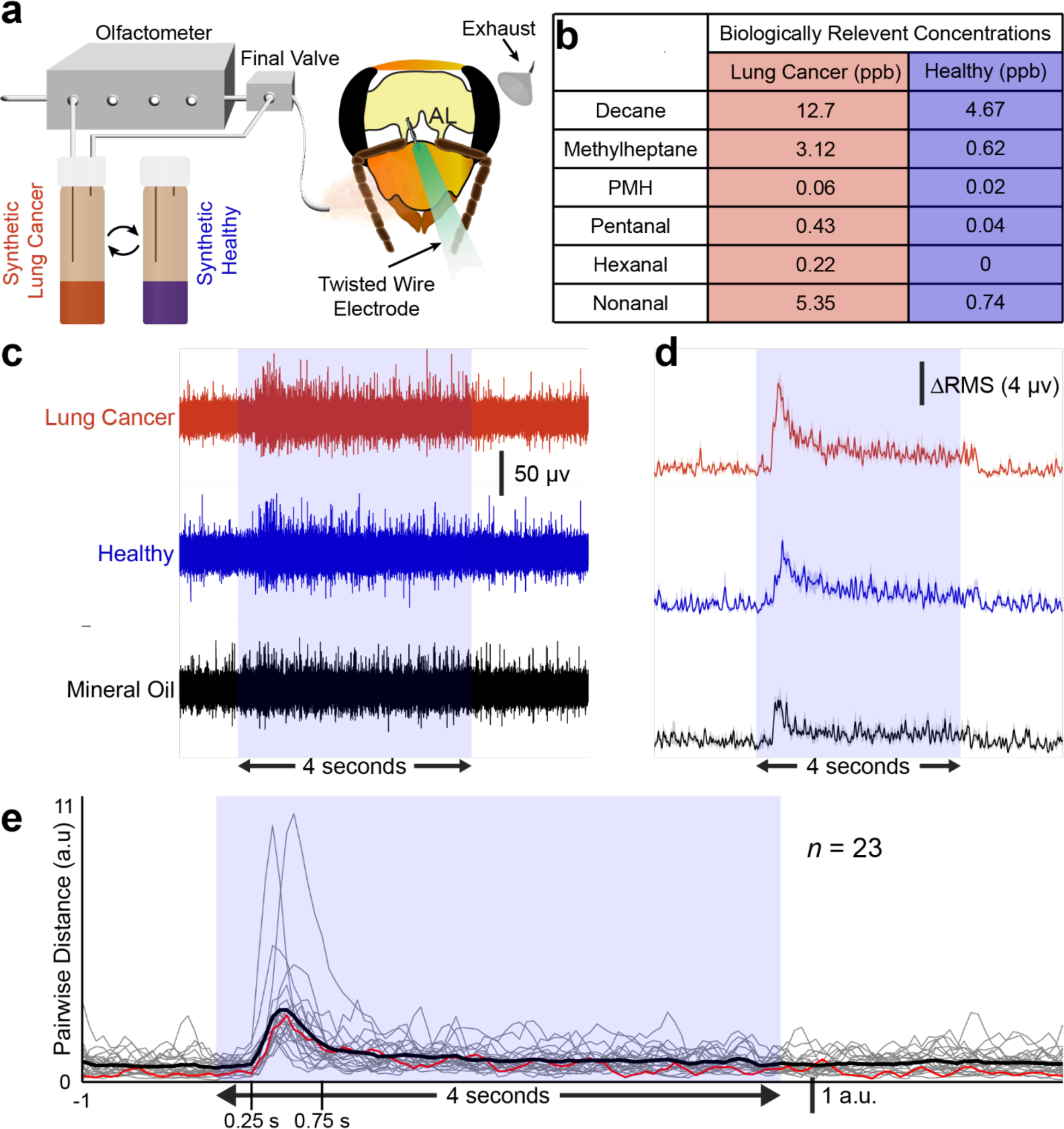
Synthetic lung cancer vs. healthy breath mixtures can be distinguished by individual ALN recordings. **(a)** Schematic of the odor delivery and neural recording setup. Notice that instead of single VOCs being delivered, now a complex mixture of VOCs (synthetic lung cancer patient breath or synthetic healthy patient breath) is delivered to the honeybee antennae. **(b)** Table of concentrations for the six VOCs mixed into the two synthetic breath mixtures. These concentrations are derived from GC-MS studies^7,88^. **(c)** Neural voltage responses of a representative ALN recording are shown for synthetic lung cancer breath, synthetic healthy breath, and mineral oil control. Notice that even at ppb level concentrations of the VOC mixtures, the neural recordings show clear odor-evoked responses during the odor presentation window. **(d)** R.M.S. filtering has been applied to the voltage traces and averaged across the four channels of the twisted wire electrode. The increase in voltage after stimulus onset followed by a slightly above baseline voltage for the remainder of the stimulus preserves the response patterns observed in the voltage traces in **panel c**. **(e)** Pairwise distance plots are shown using the same method as in Fig. 2a. Here, a total of 23 R.M.S. voltage traces are plotted (grey). The red line indicates the pairwise distances for the R.M.S. trace shown in **panel d** and the black line is the average of all 23 R.M.S. voltage traces. Again, notice that the maximum pairwise distances for most positions is between 0.25 – 0.75 seconds after stimulus onset.

Even at these low VOC concentration mixtures, we observed clear odor-evoked responses from the ALNs to the synthetic breath stimuli (**Figs. 4c, d**). Instead of employing a spike rate-based analysis, the data was processed using a root mean squared (R.M.S.) transform-based approach, which can be implemented automatically and in real time (see Experimental Section). The R.M.S. transform preserves the overall shape of the voltage trace as shown in our previous work^89^. For example, at a representative position shown in **Fig. 4d**, the synthetic lung cancer breath elicited the largest peak in the R.M.S transformed odor-evoked response while the mineral oil control elicited the smallest peak.

Next, to confirm that the recording sites contain discriminatory information corresponding to synthetic lung cancer vs. healthy breath mixtures, a pairwise distance analysis, previously used for the spike sorted individual cancer biomarker-evoked responses, was implemented for the R.M.S transformed synthetic breath responses. Again, most of the recorded positions showed differences in the neural responses to the synthetic lung cancer vs. healthy breath, with the largest difference between the responses occurring during the transient odor response window from 0.25 – 0.75 seconds (**Fig. 4e**).

### 2.5. Honeybee population neural responses classify synthetic lung cancer vs. healthy breath at a high success rate

How well can lung cancer vs. healthy breath mixtures be classified at the population ALN level? To address this question, first, population neuronal trajectories were plotted corresponding to synthetic lung cancer vs. healthy breath mixtures (see Experimental Section). Briefly, R.M.S transformed neural responses were binned into discrete 50 ms bins by averaging the voltage values within each bin and subsequently converted into odor specific population matrices, using the same procedure as previously implemented on individual cancer biomarker data (**Fig. 2b**). For this analysis, each R.M.S response was considered a separate dimension and the data was transformed using PCA to reduce from all 23 dimensions to the first three PCs for visualization (**Fig. 5a**). Within the PC subspace, all three odors (synthetic lung cancer breath mixture, synthetic healthy breath mixture, and mineral oil control) were clearly projecting in different directions away from the origin demonstrating good separation between the neural responses to all three stimuli. This result indicates that although only small variations in concentrations (i.e., ppt level) exist between healthy vs. lung cancer synthetic breath mixtures, they are encoded differently by the neuronal population in the honeybee antennal lobe, which can be leveraged to classify those mixtures using population neuronal response templates.

**Figure 5:**
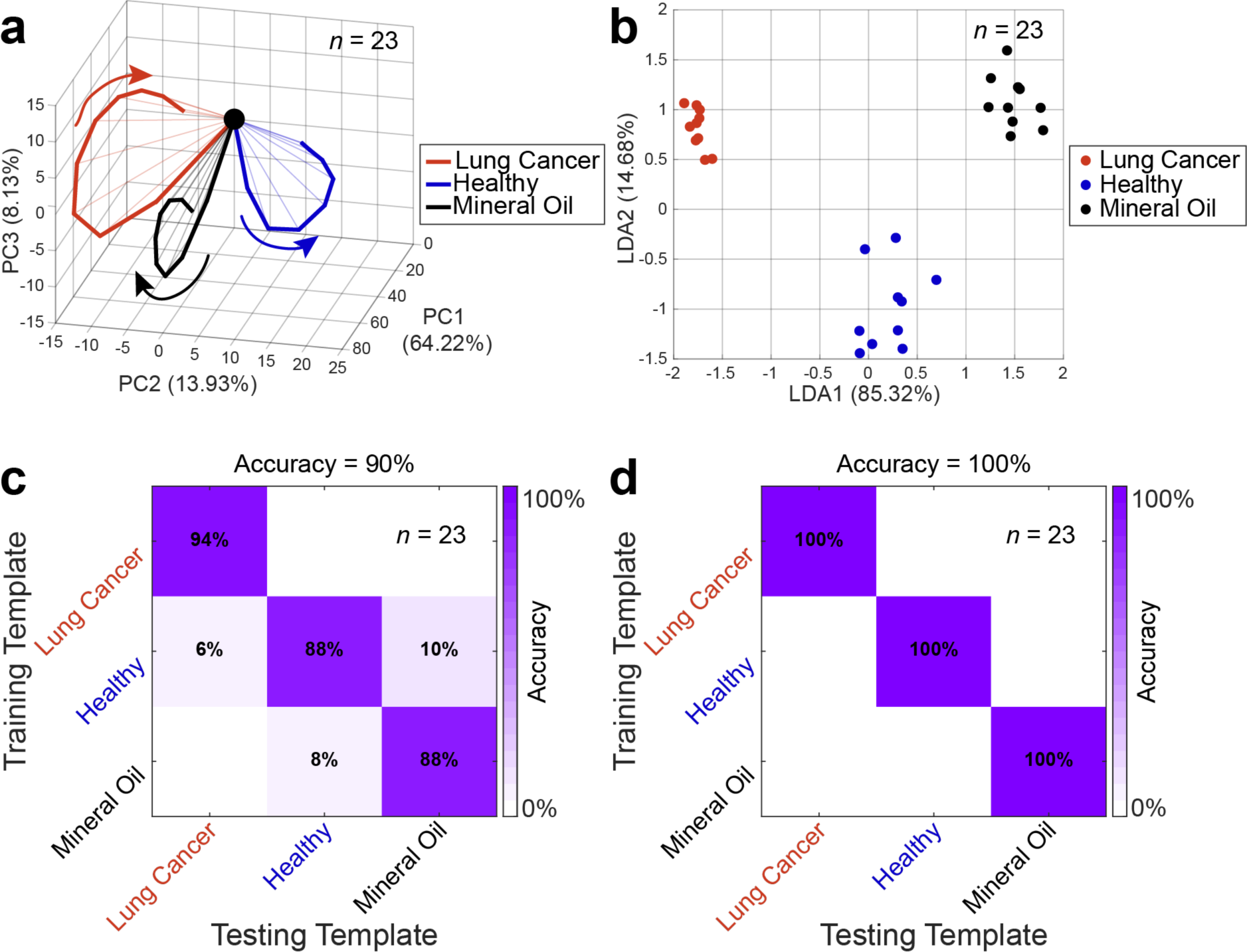
Analysis of the population neural responses classifies the synthetic lung cancer vs. healthy breath mixtures correctly. **(a)** PCA trajectories of the 23 R.M.S. voltage traces of the three odors all project in different directions. **(b)** A 3-class LDA, a supervised dimensionality reduction technique, clearly separates the odors into three distinct clusters. **(c)** LOTO analysis of each time bin, as described for Fig. 3g, correctly classifies the majority of time bins as indicated by the higher diagonal values in the confusion matrix. **(d)** LOTO analysis of the entire transient response period (0.25 – 0.75 seconds after stimulus onset) correctly classifies every single trial of all three odor mixtures tested (100% accuracy).

Next, we tested how well the lung cancer vs. healthy breath mixture-evoked neuronal responses are separated using a second technique, linear discriminant analysis (LDA). This is a supervised algorithm that maximizes between-cluster separation while minimizing within-cluster variance. We observed that the three odors formed separate, distinct, and cohesive clusters (**Fig. 5b**) indicating good separation between them.

Finally, to quantify the classification success rates of lung cancer vs. healthy breath mixtures, we employed a high dimensional LOTO analysis (see Experimental Section). For time bin-wise classification, 90% of the total bins were classified correctly (**Fig. 5c**). When we looked at the classification success across the entire 0.25 – 0.75 second time window, which we already established as the most discriminatory part of the neural signal, the classification success reached 100% using the LOTO approach (**Fig. 5d**). We performed the same analysis using spike sorted ALN data on the synthetic breath mixtures and obtained similar classification results (**Supplementary Fig. 2**).

### 2.6. A simple firing rate-based analysis is enough to classify lung cancer vs. healthy synthetic breath from separate training and testing data sets

As the LOTO approach yielded near perfect classification, we asked: how well can ALNs classify lung cancer vs. healthy synthetic breath when the training and testing datasets are completely separate? To test this, the three target odors (synthetic lung cancer, synthetic healthy breath, and mineral oil) were initially presented in a pseudorandom order for five trials each. Following a 10-minute break during which only clean air was presented to the honeybee antennae, the same three odors were presented for another five trials each in a new pseudorandom order. Using this odor presentation procedure, the first five trials of each odor were used to form the training templates, while the last five trials were used as testing templates (**Fig. 6a**).

**Figure 6:**
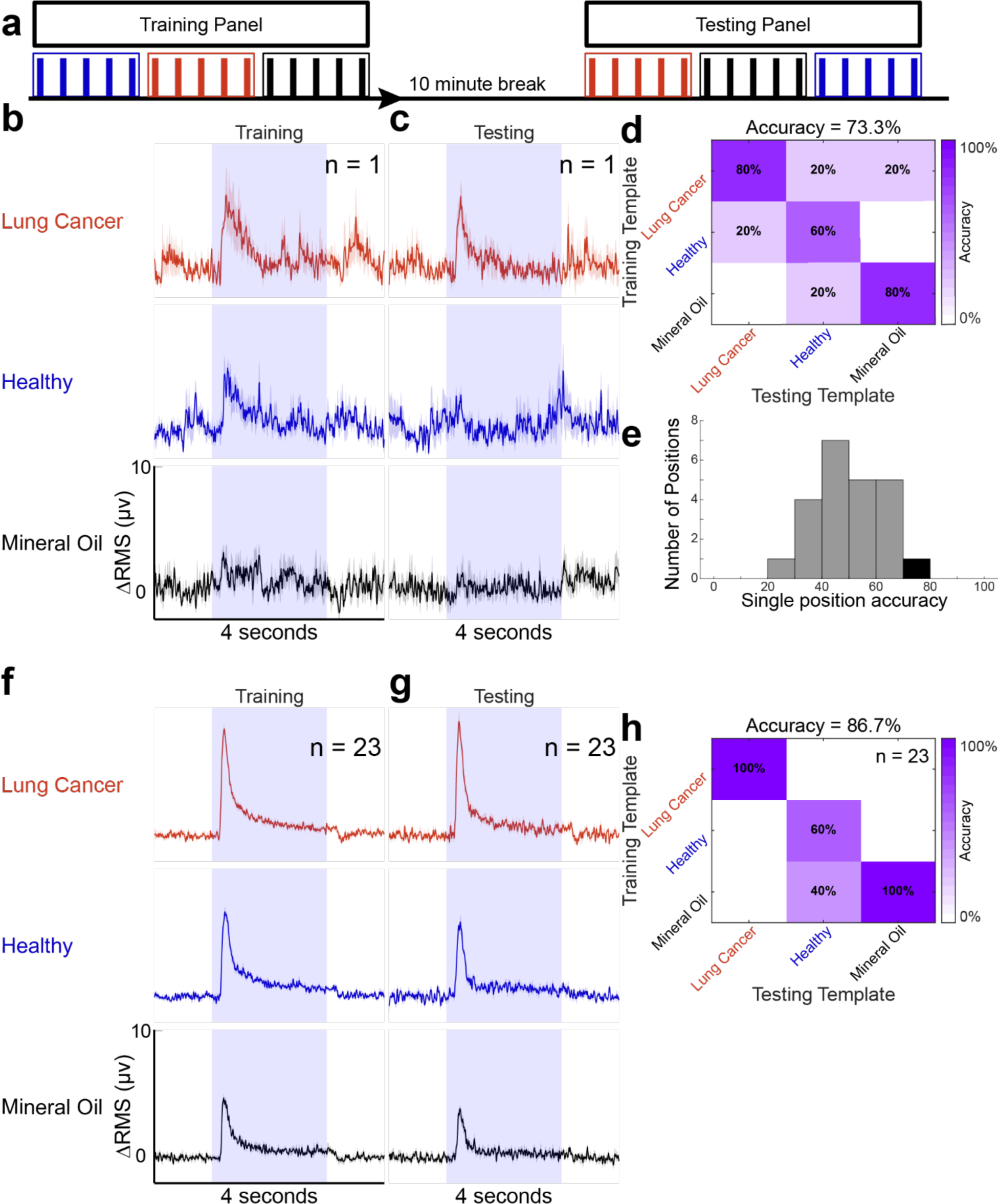
With temporally separate and independent training and testing phases, synthetic lung cancer breath vs. synthetic healthy breath mixtures can be classified using a simple population neural response motif. **(a)** Schematic demonstrating the odor stimulus presentation timeline for a single experiment is shown. Five trials with one-minute inter-stimulus intervals of each of the three odors are all initially presented to the honeybee antennae in a pseudorandom order and used as the training data set. Then, the insects were presented with clean air for 10 minutes. Finally, the same three odors are presented to the honeybees for another 5 trials with one-minute inter-stimulus intervals each in a new pseudorandom order. The second presentation of the odors was used to create the testing data set. Red: synthetic lung cancer breath, blue: synthetic healthy breath, black: mineral oil. **(b)** Average ± S.E.M. of R.M.S voltage traces using 10 ms bins of the five trials of the training odor panel for a single recording position. **(c)** Average ± S.E.M. of the R.M.S voltage traces for the testing odor panel at the same recording position. **(d)** Classification results comparing the peak voltages of the five testing trials (x-axis) to the average of the five training trials (y-axis) at this single recording position. **(e)** Histogram of the classification accuracies using the peak voltage response criteria of all 23 recording locations are shown. The average classification value corresponding to the location shown is plotted in black. **(f)** Average ± the S.E.M. (over 5 trials) of the population R.M.S. response (23 positions) using 10 ms bins for the training trials. **(g)** Average population R.M.S responses of the testing trials. **(h)** Mean classification results comparing the peaks of the population R.M.S. responses are summarized in a confusion matrix. Overall classification success rate is 86.7%.

First, we tested the classification performance of individual recording positions (n = 23 total recording sites). We observed that even for temporally segregated independent observations, synthetic cancer and healthy breath mixtures evoked consistent ALN responses (**Figs. 6b, c**). After applying the R.M.S transformation to the odor-evoked voltage traces, the peak value during the transient odor response periods (0.25 – 0.75 second post-stimulus onset) were compared between the training and testing datasets (**Fig. 6d**). For this analysis, the five training trials for each odor were averaged together to create three training templates while each of the five testing trials from all three odors were compared to the three training templates. A confusion matrix quantifying the accuracy of a single position’s classification ability is shown in **Fig. 6d**. We observed that for this recording, 11 of the 15 testing trials were classified correctly. Individual position classification accuracy had a large, unimodal distribution (**Fig 6e**), indicating that some positions encoded each odor distinctly and reproducibly across the training and testing trials thereby ensuring good classification while other positions did not have meaningful differences between odors and therefore produced low classification success. However, this is expected for any odor classification at the individual neuronal level in the antennal lobe and therefore we pooled multiple neurons/recording sites to perform population response-based classification analysis.

As it was observed at the single neuron level that the peak neural response within the transient period is a good metric for distinguishing synthetic lung cancer vs. healthy breath, we performed the same analysis combining all recording locations (n = 23). From the population R.M.S traces (**Figs. 6f, g**) we observed that the synthetic lung cancer breath had the largest peak, indicating the largest amount of neural activity occurred during synthetic lung cancer breath presentation. The synthetic healthy breath evoked a moderately sized peak while the mineral oil control had the smallest peak. By quantifying these differences using the same comparison of peaks between the training and testing templates, we achieved 100% correct classification of the synthetic lung cancer breath. Two of the five trials of the synthetic healthy breath were misclassified as the mineral oil control, and we achieved 100% correct classification of the mineral oil control (**Fig. 6h**).

## 3. Discussion

Early detection of cancer is essential to improve patient outcomes by allowing for a greater chance of successful treatment, and therefore a higher likelihood for recovery. The current recommended technology for lung cancer screening is a radiation imaging technique called low-dose commuted tomography (LDCT)^90^. Using this method to screen high-risk patients has improved overall lung cancer mortality, but this technique is not readily available to everyone, has known issues with false positives, and the associated risks due to exposure to ionizing radiation prohibit use for screening in the general population^91^. As mentioned previously, breath-based diagnostics offers a promising alternative screening modality due to its inherent noninvasiveness and the possibility for reliable and early detection of cancers. The human body emits a variety of VOCs that are released from breath, sweat, urine, feces, and vaginal secretions^92^. Different factors such as diseases^20–24^, exercise, food consumption, and drug use^93^ are known to alter the components and concentrations of VOCs, and thus can reflect the metabolic condition, or health, of an individual. A wide variety of VOCs at specific concentrations have been associated with cancer from multiple *in vitro* and *in vivo* studies spanning lung, breast, leukemia, gastric, and prostate cancer^94–103^. Different chemical compounds including hexanal, propanal, pentanal, acetone, pentane, 2-butanone, and benzene at specific concentrations have been identified as potential lung cancer biomarkers^104^. Overall, there is growing evidence that VOCs in the breath of cancer patients are altered compared to healthy individuals^105–108^, and these unique compositions of VOCs in exhaled human breath can be used as a fingerprint for the early detection of cancer. We have shown that honeybees can detect all nine of the different lung cancer associated VOCs we tested, indicating their capacity to be a breath-based diagnostic sensor.

E-noses are portable sensors that implement biological principles of gas sensing to achieve one-shot complex VOC mixture classification. They employ biological principles in detecting and quantifying VOCs in a breath sample by converting the interaction of VOCs with different materials into electrical signals, which are then processed using cross-selective sensors and template-matching analyses^66,109,110^. These e-nose devices have also been used to effectively classify multiple types of cancers, including lung^111,112^, prostate^113^, and head and neck^114^. E-noses are simpler and more portable than GC-MS, making them suitable for point-of-care and home-based VOC analysis. However, e-noses have a lower specificity and sensitivity compared to GC-MS and are generally not able to detect all the VOCs in a breath sample, especially at low concentrations. E-noses also have low-reliability over time due to sensor drift, and their responses can be altered due to changes in environmental conditions such as temperature and humidity^63,109^. Some e-noses have recently started incorporating biological olfactory receptors and live cells for VOC sensing^115–125^. However, using only one or few biological receptors does not provide robust detection capabilities for these e-noses and their reliability of detection over time remains an issue. Using the broadly sensitive honeybee olfactory sensory system, with no need to know or design the sensor for specific VOC identity or concentration ranges, we reliably classified all nine potential lung cancer biomarkers and the synthetic breath mixtures.

Biological olfaction, such as a dog’s nose, has been used to detect different types of cancer, including lung cancer, from human breath samples^69,73,126–132^. Similar to dogs, more recently, behavioral studies involving other animals including rats, honeybees, fruit flies, and ants have been conducted to detect different diseases^132–138^. However, employing behavioral outcomes to identify a disease has its own limitations. Behavioral outcomes are binary - ‘yes’ or ‘no’, which limits this approach to detect only one disease. Behavioral studies of other animals except dogs are performed in tightly controlled environments and are susceptible to behavioral variations due to other sensory cues. Our approach eliminates behavioral variability by directly recording the cancer VOC-evoked neural signals from the honeybee’s olfactory brain, which is known to be odor identity and intensity specific, and can detect odor stimuli in natural environments^85,87,139–145^. Also, insect antennae are very sensitive, with a recent study indicating the detection threshold is sub-ppb^146^. Therefore, we expect this honeybee brain-based odor detection approach will be robust, be sensitive to the minute VOC concentrations in human breath, and work efficiently in natural settings.

Our previous work demonstrates that neural recordings from the locust brain can be employed to successfully differentiate between three types of human oral cancer cell cultures from a human noncancer cell line^89^. However, in this previous work, we could not determine which VOCs were different between the oral cancer vs. noncancer cell lines as this biological neural template-based odor classification approach does not identify individual compounds. Therefore, here, we reverse-engineered the mixture of specific lung cancer VOCs that have been identified and shown to be different between lung cancer vs. healthy human breath samples through GC-MS based studies. We mixed those VOCs at precise concentrations to generate the lung cancer and healthy synthetic breath mixtures. By using this synthetic breath approach, we were able to show that these different mixtures of six lung cancer associated VOCs can be successfully differentiated by honeybee olfactory neural recordings. Ours is the first study that confirms that mixtures of multiple volatile lung cancer biomarkers at ppb to ppt levels, mimicking those biomarkers’ concentrations in exhaled human breath, can be detected by honeybee olfactory neural circuitry. This study opens the door for employing powerful honeybee olfactory neural responses for noninvasive cancer detection.

For this study, we have used neurons from multiple honeybees for VOC classification. The limitation of this approach is that multiple recording experiments must be combined for the analysis. The number of neurons recorded for each experiment can be increased by improving the multi-electrode configuration or by adjusting the placements of the electrodes within the antennal lobe. In the future, we expect to collect ∼20-30 ALNs from one single honeybee, which would enable spatiotemporal classification of odors. The principles for achieving reproducibility between different honeybee brain-based sensors will depend on the generation of ‘distinct’ neuronal response templates corresponding to the target VOC mixtures (e.g., lung cancer vs. noncancer) and not on the identity of individual neurons. As we are performing VOC-evoked extracellular neuronal recordings from the honeybee antennal lobe we expect that we will not be recording from the same neurons across different brains. Here, we have recorded from both PNs and LNs. The antennal lobe contains a dense interconnected network of roughly 800 PNs and 4000 LNs, both of which contain discriminatory odor information^147,148^. In this brain-based VOC sensing approach, the identity of individual neurons is not important, rather the properties of the entire recorded ALN population responses are the key. Our goal will be to create VOC-evoked ALN population response templates that are distinct for different odors in the neural response space from a single honeybee.

Currently, the honeybee brain-based sensor’s lifetime is limited (∼2-3 hours), but it is a high thruput device capable of running a VOC mixture test every minute (as the olfactory neurons reset to baseline firing within 60 seconds of a stimulus offset). As a proof-of-concept study to show that honeybees can detect lung cancer, we have used complex gas mixtures of six compounds at ppb to ppt levels and mixed them with clean air to replicate lung cancer and healthy human breath. However, the exact concentrations of the hundreds of VOCs in the breath of lung cancer patients and healthy individuals are unknown and therefore we cannot synthetically replicate actual human breath. In the next stage, this honeybee olfactory neural circuit-based VOC sensing approach needs to be tested on human breath samples of lung cancer patients and healthy subjects.

## 4. Experimental Section

### 4.1. Honeybee husbandry

Foraging honeybees (*Apis mellifera)* were purchased and shipped from the School of Life Sciences at Arizona State University (Tempe campus, AZ, USA). The small colonies were maintained in an incubator in 24 hours of darkness at 32°C and 60% humidity for 4 weeks at a time. For food, the bees were given 50% sucrose solution and pollen patties.

### 4.2. Honeybee surgery

The honeybees were collected individually into plastic vials, then placed briefly into ice. As soon as they stopped moving, the bees were manipulated into a plastic restraining harness with dental wax (Surgident Periphery Utility Wax) placed at the back of the head to prevent the head from moving while allowing free movement of the antennae and mouthparts. The bees were then left undisturbed on the counter for 30-60 minutes, after which they were fed 50% sucrose solution until they were completely satiated. The bees were then placed into a drawer overnight, along with a small cup of water for humidity. The next morning, the bees were tested for feeding motivation as a metric to determine their health for the electrophysiological recording. This was done by exposing the antennae to the 50% sucrose solution, and if a subsequent proboscis extension response was not observed, those bees were discarded. Eicosane wax (Sigma-Aldrich) was used to immobilize the antennae by first applying the wax to the base of the scapes and then to the pedicels, and then positioning the antennae so that the flagellum were both forward-facing. The top of the head was shaved using a microscalpel, and the bees were allowed to rest undisturbed for 10-20 minutes. Next, a small window was cut into the top of the head just above the antennae and below the ocelli. The glands and tracheae were removed to expose the brain. The brain was kept hydrated with a Ringer’s solution described by Brill et al. (in mM as follows: 37 NaCl, 2.7 KCl, 8 Na_2_HPO_4_, 1.4 KH_2_PO_4_; pH 7.2; all chemicals from Sigma-Aldrich)^149^. The antennal lobe portion of the brain was desheathed just prior to electrode placement.

### 4.3. Odor Preparation and odor stimulation

For the individual cancer biomarkers, the following odorants were used: decane, undecane, 2-methylheptane, 2,2,4,6,6-pentamethylheptane (PMH), pentanal, hexanal, nonanal, propylbenzene, and trichloroethylene (all from Sigma-Aldrich). These odors were diluted in 10 mL of mineral oil at 1% vol/vol concentration, and the pure mineral oil was used as a control.

To generate the synthetic lung cancer and healthy breath mixtures, we used the vapor pressure of each VOC and Raoult’s Law to obtain the liquid phase volumes necessary to achieve the desired gas phase concentrations of each chemical compound that have been previously reported in the literature (**Fig. 4b**)^7,88^. For the synthetic lung cancer breath mixture, the following odorants at their respective concentrations in the liquid phase were used (concentrations in μM): 1.0 pentanal, 1.1 hexanal, 390.9 nonanal, 240.0 decane, 15.2 2-methylheptane, and 0.8 PMH. For the healthy breath mixtures, the following concentrations were used (concentrations in μM): 0.1 pentanal, 0.0 hexanal, 54.1 nonanal, 88.2 decane, 3.0 2-methylheptane, and 0.3 PMH. Serial dilution was used to create these mixtures in 10 mL of mineral oil.

Odor delivery was conducted following our pre-established methodology. Briefly, a commercial olfactometer (Aurora Scientific, 220A) was used for precision odor stimulus delivery (**Fig. 1a**). At the beginning of each set of trials, 200 standard cubic centimeters per minute (sccm) of zero contaminant air was passed through the fresh air flow line via a 1/16 in. diameter PTFE stimulus flow line to the honeybee antenna, and an additional 200 sccm of zero contaminant air was passed through the dilution flow line to the exhaust. The end of the stimulus flow line was positioned approximately 2-3 mm from and pointing at the most distal antennal segment. Five seconds prior to stimulus delivery, 40% (80 sccm) of the dilution line was redirected through the odor flow line directly upstream of the odorant vials. The dilution flow and odor line joined downstream of the odorant vials, allowing for the complete mixing of the 80 sccm odor flow with the 120 sccm of the dilution flow line’s clean air. The air-volatiles mixture primed the line with volatiles up to the final valve, where the joint dilution + odor flow was directed towards the exhaust. Upon stimulus onset, the final valve redirected the fresh air flow to the exhaust and the dilution + odor flow via the stimulus flow line to the honeybee antennae. After 4 s of constant flow and stimulus delivery, the final valve redirected the fresh air flow via the stimulus flow line to the locust antenna and the dilution + odor flow back to the exhaust. One second after stimulus offset, the 80 sccm flowing through the odor flow line was redirected through the dilution flow line, mitigating any potential headspace depletion during the odor delivery. This protocol was designed to maintain a consistent flow rate through the stimulus flow line, thereby eliminating any potentially confounding neuronal responses due to mechanosensory detection of changes in air pressure. A 6′′ diameter funnel pulling a slight vacuum was placed immediately behind the honeybee during odor delivery to ensure swift removal of odorants.

### 4.4. Electrophysiology

*In vivo* extracellular neural data for the putative cancer biomarkers and synthetic breath mixtures were collected from 34 and 21 honeybees, respectively. All cancer biomarker experiments were conducted using the same stimulus panel made up of nine different VOCs and mineral oil. For the synthetic breath mixture experiments, a different panel containing synthetic lung cancer, synthetic healthy breath mixtures and mineral oil was used for all recordings. These odor panels were pseudorandomized prior to each recording, and each 4 second duration odor presentation was repeated five times with a 60 second inter-stimulus interval. Each recording session lasted 1-2 hours. Following surgery, a silver-chloride ground wire was placed into the ringer solution within the honeybee head capsule. Voltage signals from the ALNs were recorded by inserting a custom-made 4-channel electrode with impedance between 350 and 450 kΩ superficially into the center of the antennal lobe. Voltage signals were sampled at 20 kHz and then digitized using an Intan pre-amplifier board (C3334 RHD 16-channel headstage). The digitized signals were transmitted to the Intan recording controller (C3100 RHD USB interface board) prior to being visualized and stored using the Intan graphical user interface and LabView data acquisition system.

### 4.5. Spike sorting

All neural data was imported into MATLAB after high pass filtering using a 300Hz Butterworth filter. The data was analyzed by custom-written codes in MATLAB R2020b. For spike sorting analysis, all data was processed with Igor Pro using previously described methods^150^. Detection thresholds for spiking events were between 2.5 and 3.5 standard deviations (SD) of baseline fluctuations. Single neurons were identified if they passed the following criteria: cluster separation *>* 5 SD and inter-spike intervals (ISI) < 20%. For the putative lung cancer biomarker panel, a total of 44 neurons were identified from 34 honeybees. For the synthetic breath mixtures, a total of 27 neurons were identified from 21 honeybees. Spike sorted data was used for analyses in **Figs. 1, 2, 3, and Supplementary Fig. 2**.

### 4.6. R.M.S transformation of neural voltage response

After importing the data into MATLAB and processing using the 300Hz filter, for R.M.S. analysis, large and wide voltage peaks, caused by palp movements, that were greater than 15 standard deviations from the mean voltage amplitude were removed by setting the 160 samples centered around each peak (8 ms duration) equal to the mean voltage value. This was done to remove any movement artifact from the data. For synthetic breath mixture analysis, out of a total 23 positions x 6 odors (3 training + 3 testing) x 5 trials = 690 recorded R.M.S. responses, only 10 voltage traces (each with 1-2 artifacts per recording) contained this type of movement artifact. Next, the filtered data was trimmed to the time window of interest and passed through an R.M.S. filter as described in our previous work^89^. In short, a continuously moving 500-point R.M.S. filter, followed by a continuously moving 500-point averaging filter were applied to the raw voltage data. Baseline values calculated as the average of the 2 seconds prior to stimulus onset were subsequently subtracted from the data to obtain the ΔR.M.S. values. These ΔR.M.S. values were then binned into non-overlapping 10 ms or 50 ms bins and the average of each bin was computed. For the 23 recorded positions, obtained from the 21 honeybees, used for the synthetic breath mixture recordings (multiple recording positions from a single bee is possible), R.M.S. transformed voltage data from the 4-channels of the twisted wire electrode were averaged together. R.M.S transformed data was used for analyses in **Figs. 4**, **5, 6, and Supplementary Fig. 3**.

### 4.7. Pairwise distance calculation

Using the binned spike sorted or binned R.M.S data, the odor-evoked responses across all five trials for each neuron or R.M.S. position were averaged together. Then, all possible combinations of two different odors (45 combinations for 10 individual biomarkers and 3 combinations for 3 synthetic breath mixtures) were compared by calculating the absolute value of the difference between time-matched bins. The average of odor-evoked pairwise distances across all 44 neurons for the putative cancer biomarkers (**Fig. 2a**) and all 23 R.M.S. recording positions for the synthetic breath mixtures were also computed (**Fig. 4e**).

### 4.8. Dimensionality reduction analysis

We performed two methods of dimensionality reduction – Principal Component Analysis (PCA) and Linear Discriminant Analysis (LDA) as described in our previous study^89^. In PCA, we binned baseline subtracted, spike sorted neuronal signals into 50 ms non-overlapping time bins and averaged over trials (n = 5, each stimulus was repeated 5 times with a 1-minute inter-stimulus interval). The baseline response was calculated for each neuron by averaging the firing rate over the 2 second time windows immediately before stimulus presentation across trials. Recorded neurons were pooled across multiple electrophysiology experiments. For example, in **Fig. 2b**, spike sorted and binned responses of all recorded neurons (44 total) were combined to generate a *neuron number (n = 44) × time bins (t = 10, number of 50 ms bins between 250 ms – 750 ms)* matrix, where each element in the matrix corresponds to the spike count of one neuron in one 50 ms time bin. Similar neuronal population time-series data matrices were generated for each stimulus. PCA dimensionality reduction analysis was performed on the time-series data involving all 10 putative cancer biomarkers (decane, undecane, 2-methylheptane, 2,2,4,6,6-pentamethylheptane, pentanal, hexanal, nonanal, propylbenzene, trichloroethylene, and mineral oil control) and directions of maximum variance were found (**Fig. 2b**). The resultant high-dimensional vector in each time bin was projected along the principal component axes. Only three dimensions with the highest eigenvalues were considered for visualization purposes and data points in adjacent time bins were connected to generate low-dimensional neural trajectories. The same PCA analysis was applied to the synthetic breath mixtures (lung cancer, healthy, mineral oil) using the baseline subtracted R.M.S transformed population time-series data (**Fig. 5a**). For LDA analysis, the same R.M.S transformed population time-series data matrix was used. Here, we maximized the separation between interclass distances while minimizing the within class distances. To visualize the data, time bins were plotted as unique points in this transformed LDA space and stimulus-specific VOC clusters became readily apparent (**Fig. 5b**). All dimensionality reduction analyses were done using custom written MATLAB R2020b codes.

### 4.9. Quantitative classification: Leave-one-trial-out

For individual biomarker classification, the spike sorted data was binned into 50 ms bins to create a three-dimensional matrix (44 neurons *×* 5 trials *×*10 time bins). The number of time bins depends on the time window of interest, for this analysis, the time window was chosen as 0.25 – 0.75 seconds after stimulus onset, which corresponds to the transient phase of the odor-evoked neural activity that is most discriminatory. For leave-one-trial-out (LOTO) analysis, four trials (out of total five) were averaged together to form a training template, while the fifth trial, the left-out trial, was used as the testing template. For individual biomarker classification, this was done for all odors to create 10 training templates and 10 testing templates. For each time bin within the time window of interest, the Euclidean distance between the testing templates and the training templates were calculated and the testing templates were assigned based on the minimum Euclidean distance metric. Then, we iterated through the five trials, each time leaving out a different trial for the testing template and using the other four trials to create new training templates. The results are summarized with a confusion matrix (**Fig. 3g**). A fully diagonal matrix indicates 100% classification accuracy with any deviations indicating misclassifications. A further analysis was done by calculating the mode for each testing template across the 10 time bins in the time window of interest. The mode was used to classify the entire trial in a winner-take-all approach and the results are summarized in **Fig. 3h**. This same analysis was done on the R.M.S. transformed synthetic breath mixtures (**Figs. 5c, d**). Here, the three-dimensional matrix created for each odor is 23 positions *×* 5 trials *×* 10 time bins.

### 4.10. Separate Training and testing dataset analysis

By presenting the synthetic breath mixtures to the honeybees at two different temporally separated times, the data can be split into independent training and testing sets. The voltage traces were transformed into R.M.S. voltage traces with 10 ms bin sizes. The first five trials and all 23 positions of R.M.S.-transformed data were averaged together from each odor to create three training templates from the 23 positions. Each of the five trials of the R.M.S.-transformed data from each odor after the 10-minute pause were used to create 15 (5 trials x 3 odors) separate testing templates from each of the 23 positions. The testing templates were assigned to the training template that minimized the difference in peak voltage value from 0.25 – 0.75 seconds after stimulus onset. The predicted assignments were then compared to the true class labels for each testing template in a confusion matrix. A representative R.M.S. transformed position is shown in **Figs. 6b-d** and the accuracies of each individual position are summarized in a histogram (**Fig. 6e**). Then, all 23 R.M.S. transformed positions were averaged together into a population R.M.S. response (**Figs. 6f, g**) and the same comparison of peak values during the 0.25 – 0.75 seconds after stimulus onset was done (**Fig. 6h**).

## 5. Conclusions

Our study demonstrates for the first time that a powerful biological VOC sensor, the honeybee olfactory brain, can be leveraged to detect human lung cancer biomarkers and complex mixtures of biomarkers at biological concentrations. By employing *in vivo* neural recordings from the honeybee brain as a noninvasive biosensing approach for lung cancer detection, we are able to combine the power of the entire biological olfactory sensory array (sensory neurons at the honeybee antennae), biological chemical transduction, and downstream neural network computations of the antennal lobe in one single brain-based VOC sensing device. We have also demonstrated that biological neural computational analyses can be performed on the biomarker-evoked neural data to achieve high classification success for synthetic lung cancer vs. healthy breath mixtures. This novel study opens the door for more forward engineering approaches for cancer detection using honeybee olfactory neurons.

## Supporting information

Supplementary Informations

## Acknowledgements

This work was supported by an NSF CAREER grant to D.S., and Startup from Michigan State University to D.S. We thank Dr. Gene E. Robinson (UIUC) for providing feedback on initial versions of the story, Dr. Zachary Huang (MSU) for his advice on honeybee husbandry and providing the initial honeybees for protocol development, and Dr. Brian H. Smith (ASU) for advice on electrophysiology recordings and to facilitate supply of honeybees from Arizona to MSU.

## CRediT authorship contribution statement

D.S. conceptualized the study. D.S., E.C., M.P., and A.F. designed experimental plans. E.C. conducted electrophysiology experiments. Data analysis was done by M.P., S.S., A.F., N.L., S.M. The paper was written by D.S., E.C., and M.P. All the authors contributed to review and editing of the manuscript. D.S. performed overall project supervision and administration.

## Data availability

All raw datasets generated during the study are available from the corresponding author on request.

## Code availability

All analyses were performed using custom written MATLAB codes. All codes are available from the corresponding author on request.

