## Supplementary Informations for "Employing a honeybee olfactory neural circuit as a novel gas sensor for the detection of human lung cancer biomarkers"

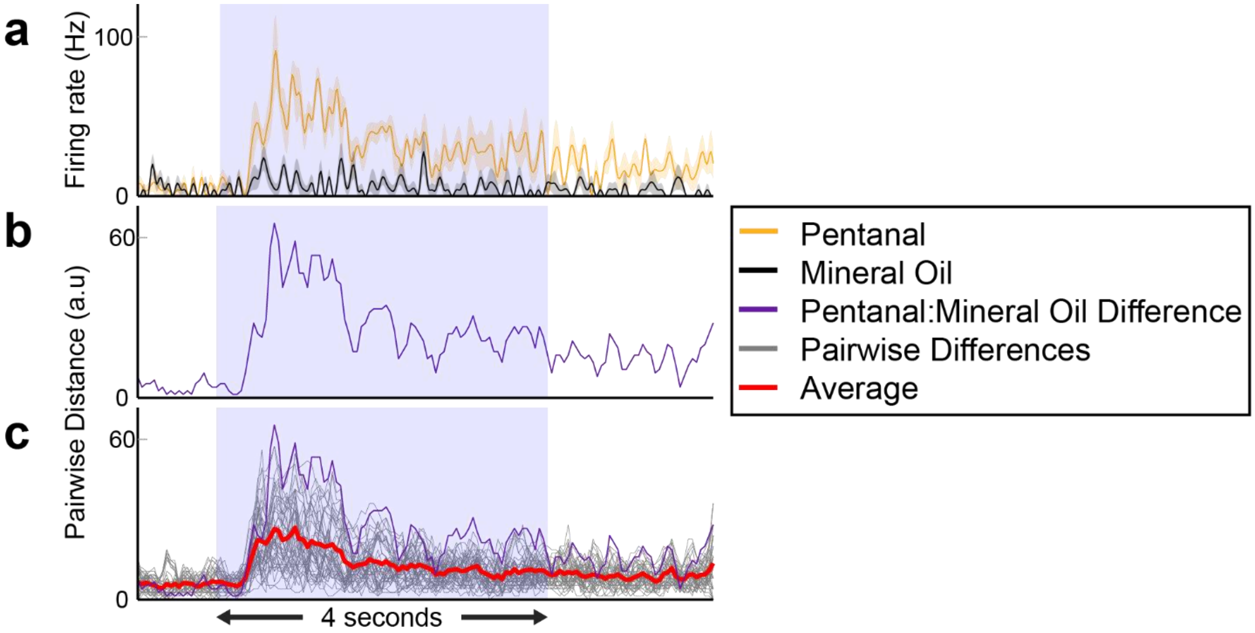


**Supplementary Figure 1: Schematic of pairwise distance analysis between odor-evoked responses. (a)** Peri-stimulus time histograms (PSTH) of a single neuron's responses to pentanal and mineral oil. **(b)** The absolute value of the difference between pentanal and mineral oil. **(c)** The grey lines are all possible pairwise differences (distances) between the ten odors (total 45 combinations) for a single neuron. The purple line is the same pentanal and mineral oil difference as shown previously. The red line is the average of all pairwise distances for this neuron. This same analysis has been done for all 44 spike sorted neurons and is plotted in **Fig. 2a**.

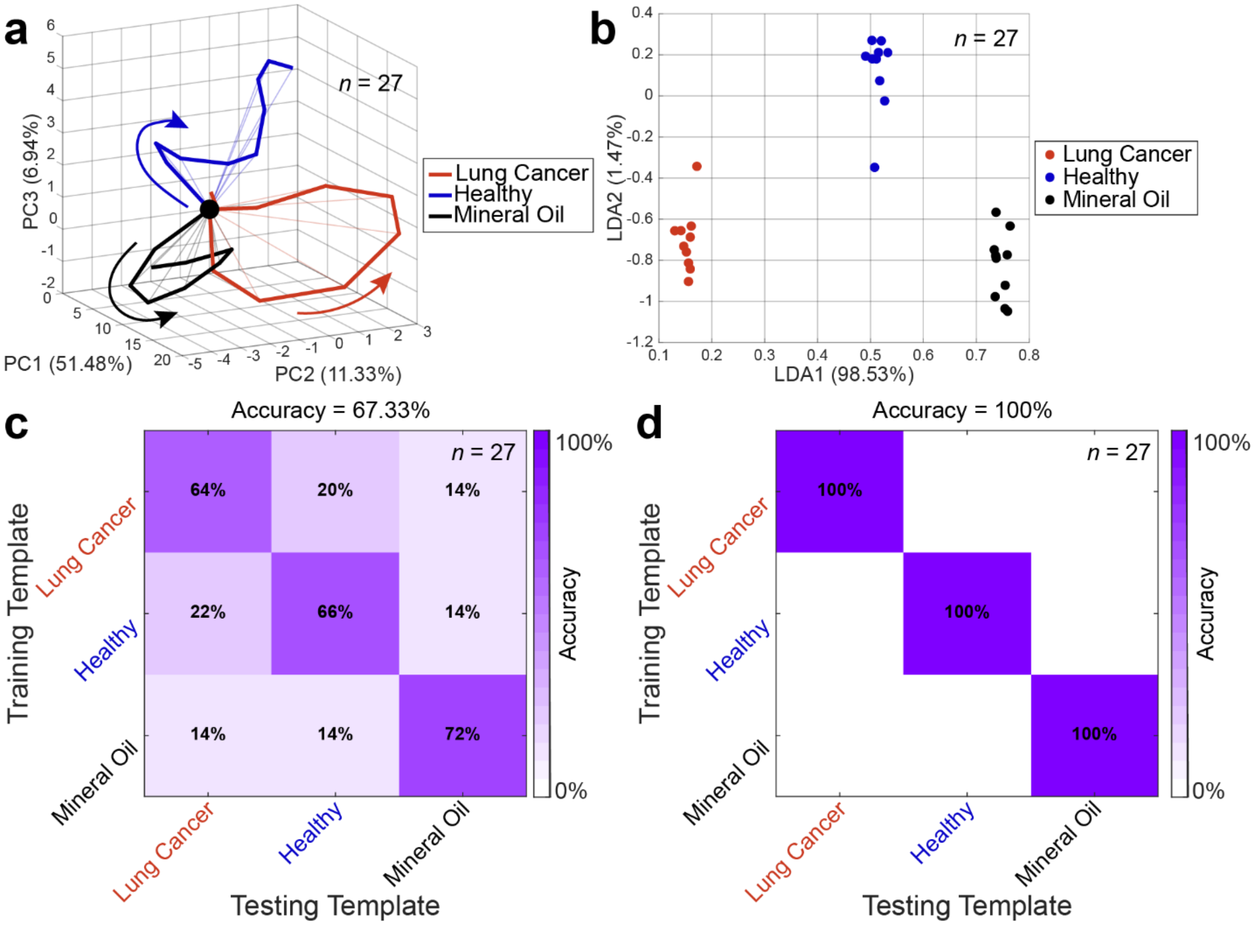


**Supplementary Figure 2: Spike sorted population neural responses classify the synthetic lung cancer breath mixture vs. healthy breath mixture correctly. (a)** PCA trajectories of the 27 spike sorted neurons of the three odor mixtures all project in different directions. **(b)** A 3-class LDA, a supervised dimensionality reduction technique, clearly separates the odors into three distinct clusters. **(c)** LOTO analysis of each time bin, as described for **Fig. 3g**, correctly classifies the majority of time bins as indicated by the higher diagonal values in the confusion matrix. **(d)** LOTO analysis of the most discriminatory response period (0.25 – 0.75 seconds after stimulus onset) correctly classifies every single trial of all three odor mixtures tested with 100% accuracy.

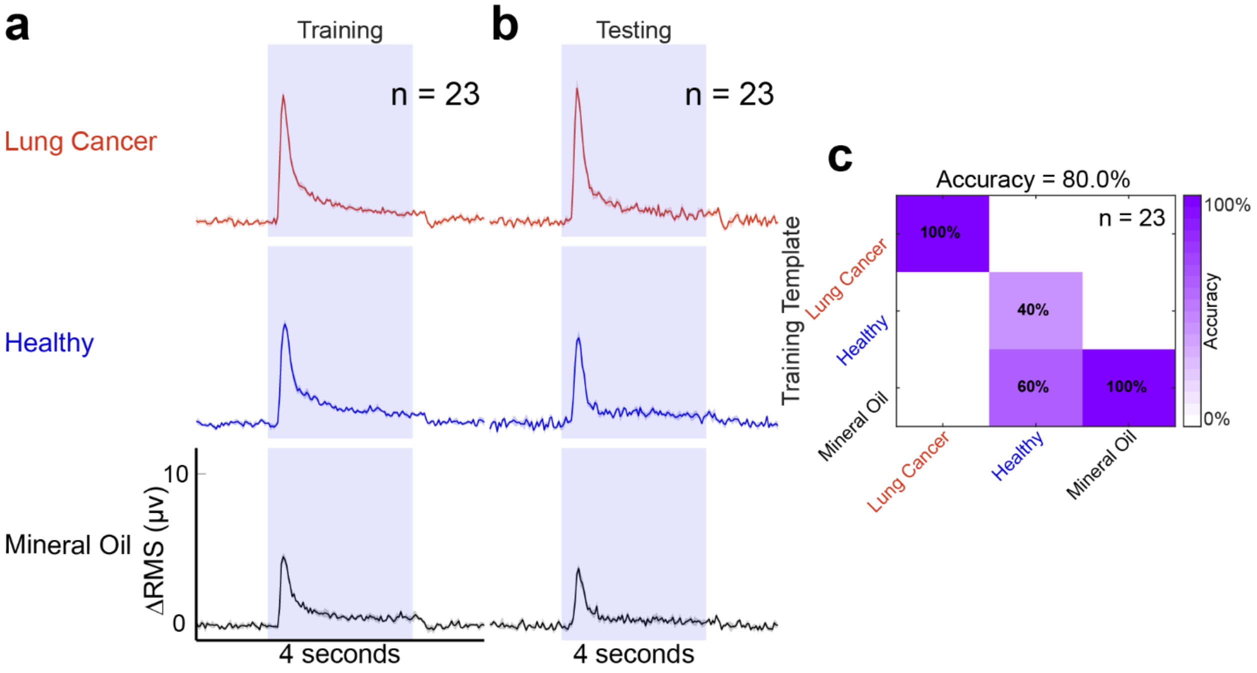


**Supplementary Figure 3: Classification of temporally separate training and testing ΔR.M.S. data is robust at different bin sizes. (a)** Average ± S.E.M. (over 5 trials) of the population R.M.S. responses (23 positions) are shown using 50 ms bins for the training trials. **(b)** Population R.M.S responses of the testing trials using 50 ms bin size. **(c)** Mean classification results are shown comparing the peaks of the population R.M.S. responses.
